# A perfusion-independent high-throughput method to isolate liver sinusoidal endothelial cells

**DOI:** 10.1101/2024.03.11.584492

**Authors:** Anna Babin-Ebell Gonçalves, Yifang Mao, Tinja Baljkas, Felix Wiedmann, Franziska Pilz, Manuel Winkler, Sina W. Kürschner-Zacharias, Marlene Hoffarth, Constanze Schmidt, Sergij Goerdt, Philipp-Sebastian Reiners-Koch, Mahak Singhal

**Affiliations:** AngioRhythms in Health and Disease, European Center for Angioscience (ECAS), Medical Faculty Mannheim, Heidelberg University, Mannheim, Germany; Faculty of Biosciences, Heidelberg University, Germany; Medical Faculty Mannheim, Heidelberg University, Mannheim, Germany; Department of Dermatology, Venereology and Allergology, University Medical Center and Medical Faculty Mannheim, Heidelberg University, and Center of Excellence in Dermatology, Mannheim, Germany; Angiodiversity and Organ Function, European Center for Angioscience (ECAS), Medical Faculty Mannheim, Heidelberg University, Mannheim, Germany; Department of Cardiology, University Medical Center and Medical Faculty Heidelberg, Heidelberg University, Heidelberg, Germany; DZHK (German Centre for Cardiovascular Research), partner site Heidelberg/Mannheim, University of Heidelberg, Heidelberg, Germany

**Keywords:** Liver, LSEC, endothelial cells, perfusion-independent

## Abstract

Liver sinusoidal endothelial cells (LSECs) critically regulate homeostatic liver function and liver pathogenesis. However, the isolation of LSECs remains a major technological bottleneck in studying molecular mechanisms governing LSEC functions. Current techniques to isolate LSECs, relying on perfusion-dependent liver digestion, are cumbersome with limited throughput. We here describe a perfusion-independent high-throughput procedure to isolate LSECs with high purity. Indifferently from previous perfusion-independent approaches, chopped liver tissue was incubated in the digestion mix for 30 minutes with intermittent mixing with a serological pipette. This led to the safeguarding of LSEC integrity and yielded 10 ± 1.0 million LSECs per adult mouse liver, which is far higher than previous perfusion-independent protocols and comparable yield to established perfusion-dependent protocols for isolating LSECs. Combining magnetic and fluorescence-activated cell sorting (FACS), LSECs from different zones of the hepatic sinusoid can now be isolated in high numbers in less than two hours for downstream applications including proteomics. Our protocol enabled the isolation of LSECs from fibrotic liver tissues from mice and healthy liver tissues from higher vertebrate species (pigs), where traditional perfusion-based digestion protocols have very limited application. In conclusion, these technical advancements reduce post-mortem changes in the LSEC state and aid in reliable investigation of LSEC functions.

## Introduction

The liver is the metabolic powerhouse of our body. While hepatocytes serve as key functional units of the liver driving major metabolic processes, LSECs have emerged as critical modulators of hepatic function in the past decade^1^. LSECs are anatomically unique as they lack an organized basement membrane and harbor fenestrae to facilitate macromolecular transport to and from hepatocytes^2,3^. Further, LSECs secrete angiocrine signals to orchestrate hepatocyte functional zonation. In line, genetic ablation of Rspondin 3, a canonical Wnt activator primarily derived from LSECs, disrupted the metabolic zonation of hepatocytes along a liver sinusoid in adult mice^4^. Likewise, angiocrine signals Wnt2 and Wnt9b regulate the expansion of Axin2+ hepatocytes to maintain hepatocyte turnover under homeostatic conditions^5,6^. Additionally, recent evidence suggests an age-related decline in LSEC function predisposes the liver towards a higher risk of developing steatosis^7^. Together, these studies highlight the gatekeeper functions of LSECs in maintaining liver health. Yet, these new findings have raised more questions about the functionality of LSECs and how can angiocrine signals be exploited as potential therapeutic targets to ameliorate liver pathologies.

Attributed to the discontinuous structure of sinusoidal vasculature, the isolation of healthy LSECs has been a challenging bottleneck for pursuing *ex vivo* functional analysis. Numerous LSEC isolation procedures have been described with the majority of those employing perfusion of the liver with tissue digestion buffer followed by a combination of magnetic and FACS-based enrichment of LSECs^8^. While perfusing the liver with digestion buffer allows isolating LSEC, it poses three major challenges – i) perfusing *per se* applies mechanical shear that potentially induces molecular alterations in LSECs’ gene circuits, ii) it strongly limits the throughput as a scientist has to perfuse one mouse at a time, and iii) it requires additional equipment and training before a scientist can insert a cannula to perfuse the liver. LSEC isolation becomes even more challenging in mice with liver pathologies including fibrosis and cancer. The accumulation of excessive extracellular matrix requires longer perfusion times with digestion mix and negatively affects the yield of LSEC isolation.

To tackle these challenges and isolate LSECs in high purity, the present study describes a perfusion-independent protocol for LSEC isolation. By opting for a perfusion-independent procedure, we circumvented the need for specialized equipment and training as well as substantially improved the scale of LSEC isolation. Focusing on the first step of the procedure – the digestion of the liver tissue, we found that intermittent mixing of tissue suspension with a serological pipette, instead of a syringe-needle combination, safeguarded LSECs and enabled a yield that is comparable or higher to previously reported yields from perfusion-dependent isolation protocols^8^. Further characterization revealed that our isolation procedure led to a uniform capturing of LSECs from all zones of the hepatic sinusoid, allowing for zone-specific LSEC isolations. Together, the newly established LSEC isolation protocol enables life scientists to study liver vasculature with minimal prior knowledge and training and propels further research efforts in understanding LSEC biology.

## Results

### Step-by-step procedure for isolating liver sinusoidal endothelial cells

The whole procedure can be classified into three major steps (**Fig. 1**):

i. Liver tissue processing to obtain single-cell suspension (∼45 mins)
ii. Enrichment of non-parenchymal cells (∼30 mins)
iii. Isolation of high-purity LSECs (∼40-60 mins)

**Figure 1.**
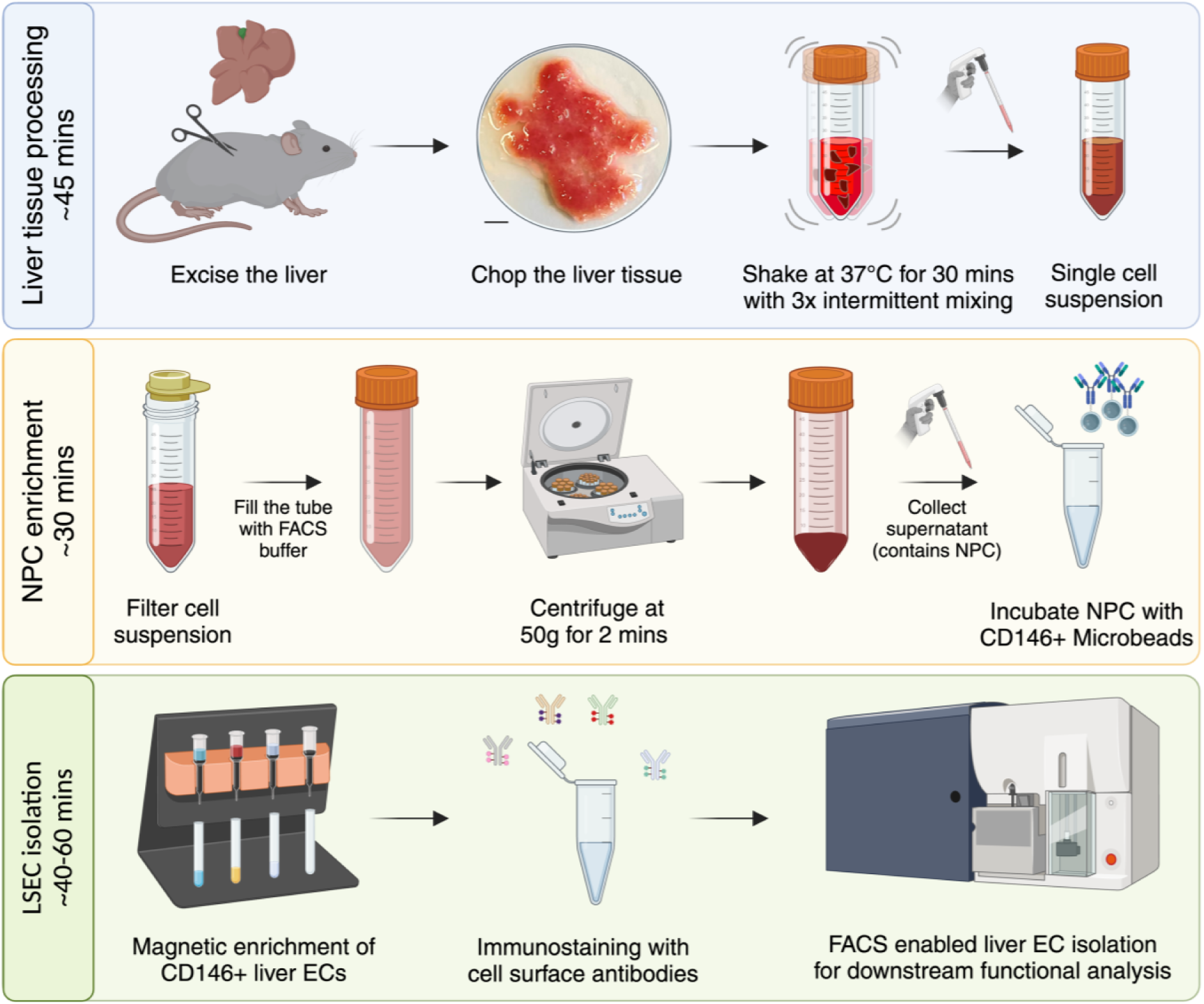
Step-by-step schematic depiction of the LSEC isolation protocol. Scale bar = 5 mm.

△ The time required for the isolation depends on the number of mice processed in parallel as well as the total cells sorted via FACS. Additionally, it would rely on FACS settings used for cell sorting. FACS sorting with a 70 μm nozzle allows for high-speed analysis with up to 15,000 events processed per second, while a 100 μm nozzle would allow cell sorting with a speed of up to 10,000 events per second. The selection of FACS settings would largely depend on the downstream application of isolated LSECs. The time indicated here is based on processing four mice in parallel and isolating 1 million LSECs for gene expression analysis.

### Step I - Liver tissue processing

1. Before sacrificing the mice, freshly prepare the digestion buffer and store it at 4°C till use. → 5 ml of digestion buffer is required to digest one adult mouse liver, approximately weighing 1 gram.
2. Excise the liver and transfer it into a 6 cm petri dish containing phosphate buffered saline (PBS) solution.
3. Swiftly wash the liver in the PBS solution and transfer it to the dry lid for chopping.
4. Use curved scissors to fine chop the liver as shown in **figure 1**. △ Coarse chopping the mouse liver tissue into small fragments will result in comparable yield of LSECs as finely chopped tissue (Fig. 2C). However, it may affect intermediate steps involving 100 µm cell strainer and occasionally clog the LS column.
5. Transfer the minced liver into a 15 ml conical tube and add 5 ml of the digestion buffer.
6. Incubate the samples at 37°C in a pre-heated incubation shaker at 180 rotations per minute for 30 minutes.
7. During the digestion, perform intermittent mixing of samples with a 10 ml serological pipette every 10 minutes for a total of three times. △ This protocol was optimized using 10 ml serological pipettes with an opening mouth diameter of roughly 1.2 mm. Using a serological pipette with a different opening diameter might affect the quality of cell suspension and the overall LSEC yield. △ After 30 minutes of digestion, there might still be remaining liver fragments, yet proceed with the next step. Elongating the digestion step might negatively affect the health of isolated LSECs.

**Figure 2.**
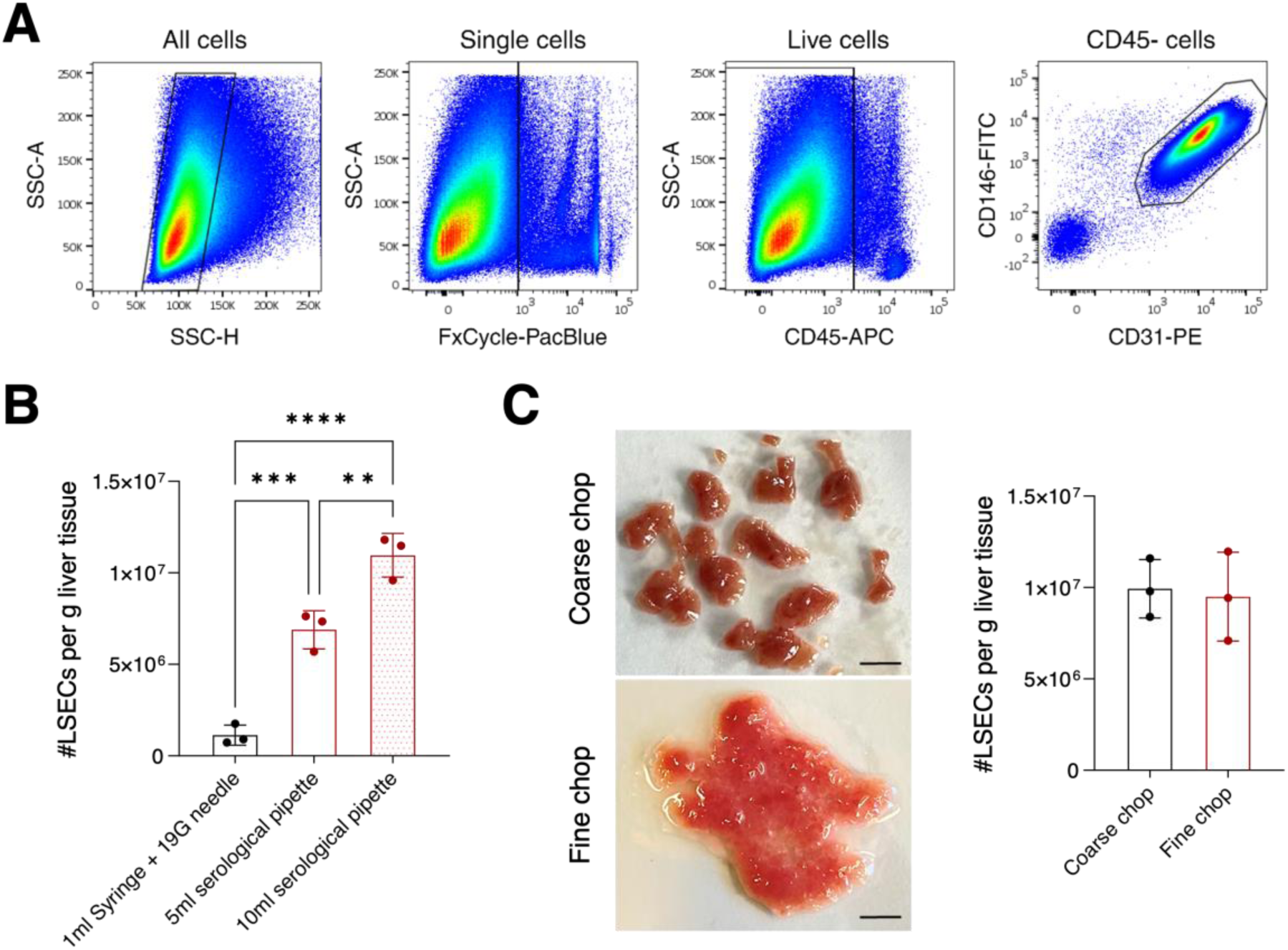
Optimizing the tissue digestion step of LSEC isolation. (**A**) Sequential FACS gating strategy to identify and sort LSECs. (**B**) The dot plot shows the count of LSECs retrieved using different methods for intermittent mixing during tissue digestion. [mean ± SD, n = 3 mice]. **P < 0.01, ***P < 0.001 and ****P < 0.0001 (one-way *ANOVA* test). (**C**) The liver tissue was either coarsely or finely chopped and used to isolate LSECs. On the left, images of differently chopped liver tissues are shown. On the right, the dot plot shows the total count of LSECs isolated from differently chopped liver tissues [mean ± SD, n = 3 mice].

### Step II – Enrichment of non-parenchymal cells (NPCs)

8. Transfer the digested cell suspension with a 10 ml serological pipette into a 50 ml conical tube through a 100 µm cell strainer.
9. In the case of a blocked cell strainer, use the plunger of a 5 ml syringe to press the remaining liver pieces through the cell strainer.
10. Wash the plunger and the cell strainer with 5 ml of FACS buffer.
11. Fill the conical tube with FACS buffer up to the mark of 50 ml.
12. Centrifuge the tube at 50g, 4°C for 2 minutes.
13. The pellet primarily contains hepatocytes, while the supernatant is enriched for NPCs. Carefully transfer the supernatant into a new 50 ml conical tube.
14. Centrifuge the tube at 450g, 4°C for 5 minutes and discard the supernatant.
15. The resulting pellet consists of NPCs. Resuspend the pellet in 200 µl of CD146-selection buffer and incubate at 4°C for 20 minutes.
16. During the incubation time, prepare for the magnetic enrichment.

a. Place the LS columns in the QuadroMACS^TM^ magnetic separator rack.
b. Equilibrate each column by adding 5 ml of FACS buffer.

### Step III – Isolation of high-purity LSECs

17. Wash the cells with 5 ml of FACS buffer and centrifuge at 450g, 4°C for 5 minutes. Subsequently, discard the supernatant.
18. Resuspend the NPC pellet in 1 ml of FACS buffer and apply it to an equilibrated LS column through a 100 µm cell strainer.
19. Wash the filter with 1 ml of FACS buffer.
20. Allow the NPC suspension to pass through the LS column.
21. Wash the LS column by applying 9 ml of FACS buffer.
22. Remove the LS column from the magnet and place it onto a fresh 15 ml conical tube.
23. Add 5 ml of FACS buffer to the LS column. Elute the cells by purging the column with the supplied plunger.
24. Centrifuge the elute at 450g, 4°C for 5 minutes and discard the supernatant.
25. Resuspend the pellet in 200 µl of the staining mix and transfer it to a 1.5 ml tube.
26. Incubate the cell suspension at 4°C for 20 minutes, protected from the light.
27. Wash the cells with 1 ml of FACS buffer and centrifuge at 450g, 4°C for 5 minutes.
28. Carefully remove the supernatant and resuspend the pellet in 200 µl of FACS buffer.
29. Transfer the sample into a FACS tube through the 35 µm cell strainer in the lid.
30. Store filtered samples in FACS tubes on ice and proceed with FACS-based cell sorting.
  a. Before analyzing a sample, add dead cell exclusion dye (FxCycle violet stain, 1 µl per sample).
  b. Sort viable CD31+CD146+ cells using the FACS gating strategy shown in **figure 2A**.
  c. Reanalyze a small portion of FACS-sorted LSECs to evaluate the purity of sorted cells.
  d. Depending on the downstream application, process the FACS-sorted LSECs.
  △ We recommend plotting a histogram of CD117/Kit as a control for successful isolation of LSECs from all regions of a hepatic sinusoid. A successful LSEC isolation will result in a plateau-like distribution in the histogram for CD117 staining. Refer to the **figure 5A**.

### Optimization of the liver tissue digestion procedure

Focusing on the tissue digestion step of the LSEC isolation procedure, we first compared different ways of intermittent mixing during incubation with the digestion mix. Here, a 19G needle - 1 ml syringe combination was compared to a serological pipette (5 ml, mouth diameter 2.5 mm or 10 ml, mouth diameter 1.2 mm) for mixing the tissue slurry. Unexpectedly, serological pipette-mediated mixing safeguarded the integrity of LSECs and resulted in a far higher yield of LSECs as compared to needle-syringe mediated mixing (**Fig. 2B**). Among serological pipettes, the use of a 10 ml pipette for mixing led to a lower chance of remaining tissue fragments after digestion on the cell strainer and a more uniform cell suspension as compared to 5 ml pipette. Subsequently, liver tissues digested with a 10 ml pipette resulted in a significantly higher number of LSECs over a 5 ml pipette (**Fig. 2B**).

Next, we employed different grades of chopping of the liver tissues, comparing coarsely and finely chopped liver tissue for LSEC isolation (**Fig. 2C**). Surprisingly, no major difference was observed in the final yield of retrieved LSECs from either of the compared chopping methods (**Fig. 2C**). However, coarsely chopped tissue often required additional effort while processing different downstream steps of the protocol. First, coarsely chopped tissue often led to incomplete tissue digestion and accumulation of leftover fragments on the cell strainer. Second, we observed a higher frequency of clogging in LS columns when enriching LSECs from coarsely chopped liver tissues. Together, these experiments suggest that finely chopped liver tissue processed with a 10 ml serological pipette offers an optimal way to digest the liver tissue for isolating LSECs.

### Isolating LSECs from different lobes of the mouse liver

Employing the above-described perfusion-independent procedure, we reliably isolated an average of 10 X 10^6^ LSECs per 1 gram of the mouse liver (**Fig. 2B-C**). Next, we compared whether our protocol could uniformly isolate LSECs from all lobes of the liver tissue. To this end, the liver tissue was segregated into three parts – left lobe, median lobes and right lobes, each constituting nearly one-third of the total liver mass (**Fig. S1A**). After that, LSECs were isolated from each of these liver parts using our protocol. We could successfully isolate LSECs from all three parts of the liver, with a yield of nearly 4 X 10^6^ LSECs from each of the segregated liver parts (**Fig. S1A**). Given that we could isolate a similar number of LSECs from three regions of the liver that weigh roughly the same suggests a broad application of the protocol to preclinical surgical experiments where often only some part of the liver is accessible for analytical experiments. A particular application would be two-third partial hepatectomy, which represents one of the well-studied tissue regeneration models^9,10^. Our method will allow scientists to isolate LSECs from the resected lobe of the liver during hepatectomy and compare them with LSECs isolated from the regenerated lobes in an indexed manner. This is often not possible with current perfusion-dependent LSEC isolation methods. This would substantially improve the robustness of lineage tracing experiments, where currently such indexed analyses are often not possible.

### Extracting LSECs from fibrotic mouse liver

Next, we questioned whether our method could be applied for isolating LSECs from fibrotic liver tissues. Accumulation of the extracellular matrix presents a technical hurdle in digesting the tissue and current perfusion-dependent isolation methods often have limited efficacy. To this end, we employed a diet-based preclinical model of metabolic dysfunction-associated steatohepatitis (MASH)^11^. Mice were fed with either standard or Choline deficient L-amino acid defined (CDAA) diet for 10 weeks to induce MASH as demonstrated by hepatocyte ballooning and accumulation of extracellular matrix in liver tissues (**Fig. 3A**). Using our method, we successfully isolated LSECs from fibrotic liver tissues (**Fig. 3B**). Further, quantitative PCR analysis revealed that LSECs from fibrotic livers have reduced expression of sinusoidal marker genes, such as *Stab1, Stab2 and Fcgrb,* as compared to LSECs isolated from healthy livers (**Fig. 3C**), suggesting the loss of sinusoidal characteristics of LSECs during fibrosis.

**Figure 3.**
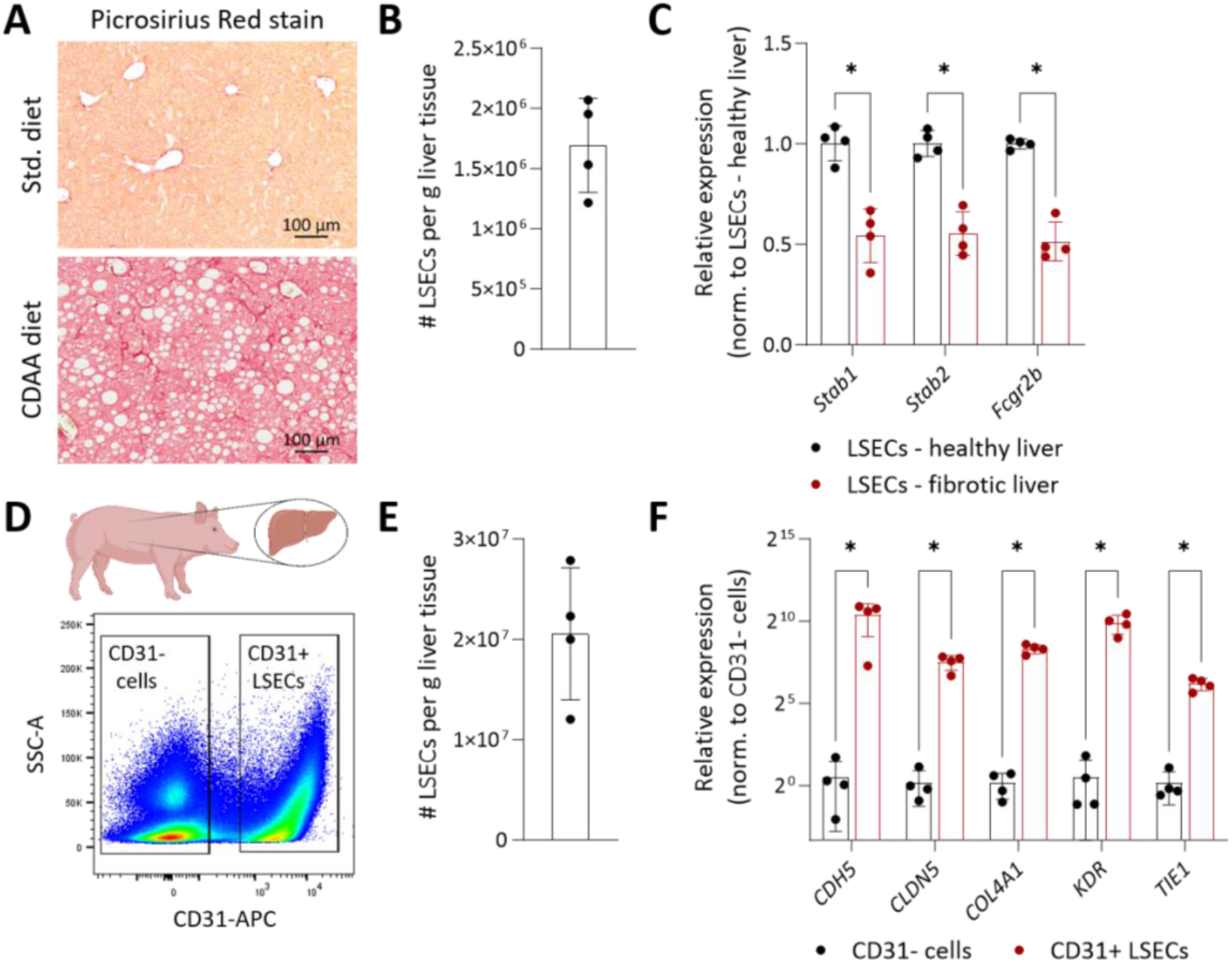
Isolating LSECs from fibrotic mouse and steady-state pig liver tissues. (**A-C**) Mice were fed with a standard or CDAA diet for 10 weeks. (**A**) Images show Picrosirius Red staining on the liver sections. LSECs were isolated from healthy and fibrotic liver tissues. Scale bar = 100 µm. (**B**) The dot plot shows the number of LSECs isolated from fibrotic mouse liver. [mean ± SD, n = 4 mice]. (**C**) Quantitative PCR analyses to compare the expression level of various sinusoidal genes in LSECs isolated from either healthy or fibrotic mouse liver tissue. [mean ± SD, n = 4 mice]. *P < 0.05 (Mann-Whitney test). (**D-F**) Liver tissues collected from healthy pigs were processed to isolate LSECs. (**D**) FACS plot showing gating strategy to isolate CD31+ LSECs and CD31-cells. (**E**) The dot plot shows the number of LSECs isolated from pig liver tissues. [mean ± SD, n = 4 pigs]. (**F**) Quantitative PCR analyses to compare expression of various vascular genes between CD31+ LSECs and CD31-cells. [mean ± SD, n = 4 pigs]. *P < 0.05 (Mann-Whitney test).

### Obtaining LSECs from pig liver tissues

To test the wider applicability of our method for isolating LSECs from the liver tissue of higher vertebrate species, we collected fresh liver samples from Landrace pigs. Processing pig liver tissues posed specific technical challenges due to the non-availability of magnetic beads for positive selection and the limited availability of fluorescence-conjugated anti-pig antibodies for FACS-based enrichment. Hence, we relied only on FACS-based enrichments with CD31 as a primary antigen to sort pig liver ECs (**Fig. 3D**) and managed to retrieve CD31+ LSECs in high numbers (**Fig. 3E**). Downstream quantitative PCR analysis illustrated the high purity of our isolated CD31+ LSECs as they manifested multi-log fold higher expression of the key endothelial genes as compared to CD31- cells (**Fig. 3F**). Relying on our isolation from pig liver tissues, we envision a broad application of our perfusion-independent protocol for isolating LSECs from small biopsies and limitedly available human liver tissues.

### Establishing the primary culture of enriched LSECs

To broaden the downstream application of our LSEC isolation method, we investigated whether LSECs can be cultured for *ex vivo* analyses. To this end, LSECs were enriched using the CD146-selection magnetic beads and then placed in a culture dish. Intriguingly, the majority of LSECs adhered to the Petri dish, suggesting the high viability of our isolated LSECs (**Fig. 4A**). Further, we interrogated whether cultured ECs retained their LSEC characteristics. To address this question, LSECs were seeded on coverslips and immuno-fluorescence stainings were performed for CD31 and CD32b/Fcγrb. While nearly all cells were CD31-positive suggesting high-purity of isolation after magnetic enrichment (**Fig. 4B**), a large fraction of cultured LSECs were double positive for CD31 and CD32b supporting the sinusoidal features of LSECs in our primary cultures (**Fig. 4B**).

**Figure 4.**
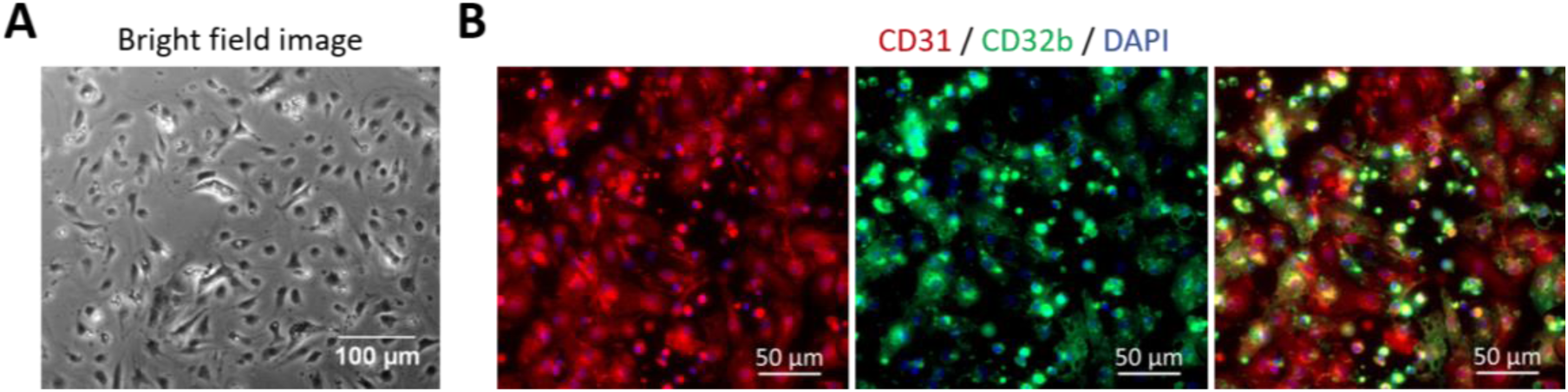
Culturing and characterizing isolated LSECs. LSECs were isolated using perfusion-independent digestion method, followed by positive selection using CD146-magnetic beads. After that, enriched LSECs were placed in culture. (**A**) Brightfield image of cultured LSECs. Scale bar = 100 µm. (**B**) Immunofluorescence images show cultured LSECs stained for CD31 (in red), CD32b/Fcγrb (in green), and DAPI (in blue). Scale bar = 50 µm.

### Retrieving LSECs from different zones of the hepatic sinusoid

Considering LSECs constitute a rather heterogeneous cell population with their function displaying a zonation along the hepatic sinusoid, we next wanted to investigate whether our protocol is capturing LSECs from all different regions of the hepatic sinusoid. To achieve this, CD31+CD146+ LSECs were sequentially analyzed for the surface abundance of CD117 (*Kit*). In line with previous publication^12^, CD117 staining on LSECs manifested a plateau-like shape (**Fig. 5A**), suggesting that our protocol was able to capture LSECs from all zones of the hepatic sinusoid. To verify the spatial origin of LSECs, we stratified and isolated LSECs based on the mean fluorescence intensity of CD117 – periportal (PP, CD117^low^), midlobular (MD, CD117^med^), and pericentral (PC, CD117^high^). Gene expression analyses corroborated the origin of sorted LSECs with periportal LSECs enriched for *Dll4, Efnb2, Il1a,* and *Sox17* whereas pericentral LSECs enriched for *Dkk3, Kit, Thbd,* and *Wnt9b* (**Fig. 5B**). Together, these data substantiate that our protocol can successfully capture LSECs from all zones of the hepatic sinusoid and present a strategy for isolating zone-specific LSECs for downstream functional analyses.

**Figure 5.**
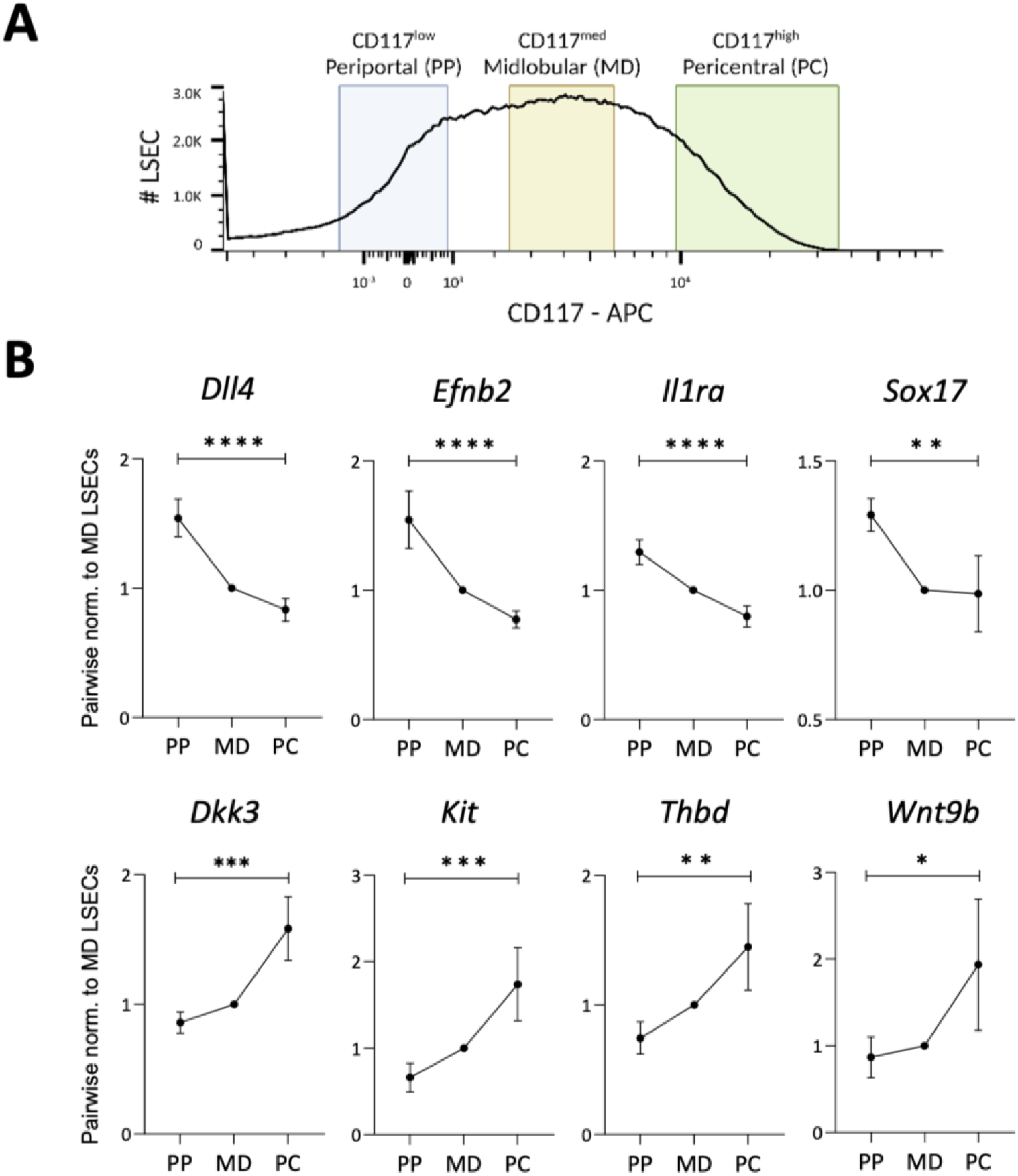
Isolation of LSECs from different regions of a liver sinusoid. (**A**) CD117-based stratification to undertake hepatic zone-specific LSEC isolation. (**B**) Gene expression analysis for known periportal (top) and pericentral (bottom) zone-specific LSEC genes [mean ± SD, n = 4 mice]. *P < 0.05, **P < 0.01, ***P < 0.001 and ****P < 0.0001 (one-way *ANOVA* test).

## Discussion

Revisiting every step of the liver sinusoidal endothelial cell isolation procedure, the present study established a perfusion-independent high-throughput method to isolate ultrapure LSECs from the mouse liver. Combining our newly established liver digestion procedure with magnetic and FACS-based purification steps, ultrapure (>98%) LSECs were retrieved in a yield far higher than existing perfusion-independent methods and a comparable yield to existing perfusion-dependent methods. Circumventing the need for perfusion means that the current protocol (i) offers scalability, allowing researchers to perform parallel isolation of LSECs from relatively large cohorts of mice, (ii) bypasses the need for substantial hands-on training to insert a cannula for perfusing the liver, and (iii) minimizes the alterations in molecular circuits of isolated LSECs driven by perfusion-induced shear stress.

### Tissue processing

Focusing particularly on the liver digestion step, we optimized the protocol to maximize tissue digestion along with safeguarding the health and integrity of LSECs. Surprisingly, use of serological pipettes instead of a needle-syringe based intermittent mixing of tissue digestion mix was important to prevent the loss of LSECs and obtain uniform results. Potentially, mixing with serological pipette reduced mechanical stress caused by syringe-needle combination, thereby preserving LSEC health. Furthermore, fine chopping of liver tissue led to smooth processing and a high recovery of viable LSECs. These two alterations in the processing of liver tissues yielded a nearly 5-fold higher number of LSECs as compared to previous perfusion-independent isolation approaches^13^.

### Digestion buffer

In the present study, we used research-grade liberase enzyme for digesting the liver tissue. This was primarily driven by the user experience as liberase offered very high consistency in terms of tissue digestion among various production lots. In contrast, the efficacy of collagenase and dispase varied depending on their lots, often requiring different concentrations of the enzymes to be used for tissue processing. Yet, all collagenase-based enzymatic digestion mixtures should work with the current protocol. We have recommended the liberase-based digestion buffer largely due to our own experiences.

### Isolating other hepatic cells

While the present study focused on optimizing the isolation of LSECs from the liver tissue, we can envision a rather broad application of the current protocol for the isolation of various hepatic cells. For example, in the step with a 50g centrifugation during NPC enrichment, the cell pellet largely consists of hepatocytes and can potentially be used for various hepatocyte-based assays. Likewise, the total NPC fraction can be used to isolate tissue infiltrating immune cells and hepatic stellate cells. While immune cells can be purified employing either cell-specific surface markers for FACS or selective adhesion for Kupffer cells^14,15^, NPCs can be processed through a Nycodenz gradient to enrich stellate cells as described previously^16^. In line, we observed Kupffer cells, marked by CD45+F4/80+ population, in our non-enriched NPCs (**Fig. S2A-B**), which could be sorted for further downstream analyses. Together, by achieving uniform digestion of the liver tissue in a perfusion-independent manner, we believe that we overcame the critical bottleneck, and our established protocol can be applied to potentially isolate all hepatic cell types.

### Broad applicability for LSEC isolation

Beyond establishing LSEC isolation from healthy mouse livers, we tested our method to obtain viable LSECs from livers where perfusion is often not possible. We demonstrate that our method efficiently retrieved LSECs from surgically resected individual lobes of the liver as well as fibrotic livers derived from MASH mice. Intriguingly, the high efficiency and purity of LSECs isolated from pig liver tissues showcase the robustness of our method for investigating LSECs from liver specimens derived from higher vertebrates and even limited clinical human material. Finally, the successful realization of primary cultures of isolated LSECs with sinusoidal characteristics makes it possible to undertake a wide array of *in vitro* mechanistic assays to study organotypic molecular mechanisms in the liver vasculature.

### Zone-specific LSEC isolation

Addressing the functional heterogeneity in LSECs across the hepatic sinusoid, we tested the surface presentation of CD117 (Kit) among CD31+CD146+ LSECs. Indeed, our isolation protocol successfully captured LSECs from all regions of the hepatic sinusoid as demonstrated by the expected trend of the expression of periportal and pericentral LSEC-specific genes. These periportal and pericentral marker genes were selected based on published single-cell RNA sequencing study of the liver vasculature^17^. Building on these findings, our method can be employed to isolate hepatic zone-specific LSECs in high quantities. In our analysis, we stratified LSECs based on the mean fluorescence intensity of CD117. Based on this parameter, the lowest 15% of LSECs were sorted as CD117^low^, and the highest 15% of LSECs were categorized as CD117^high^. This would imply that quantitatively, we could isolate around 1.5 million LSECs each from the periportal and pericentral regions of the hepatic sinusoid. The high quantity of LSECs isolated from different hepatic zones will empower future studies to undertake functional analyses such as proteomics and even observe post-translational modifications in zonated homeostatic LSECs.

In conclusion, the method described in the present study should facilitate high-throughput isolation of ultrapure LSECs. The perfusion-independent nature of the procedure should ensure high reproducibility of experiments and favor a broad application of the isolation method in laboratories that classically have little liver expertise. The current experiments demonstrate our continuous efforts to simplify the isolation protocol for different hepatic cells to better understand cellular and molecular crosstalk within the liver microenvironment.

## Acknowledgments

The authors sincerely thank Dr. Ki Hong Lee for his critical feedback on the manuscript. We thank Dr. Ashik Ahmed Abdul Pari for his help with the generation of figures with www.BioRender.com. Additionally, we are most grateful for the excellent technical support of the FlowCore and Animal Core facilities of the Medical Faculty Mannheim of Heidelberg University.

## Author contributions

ABEG, YM, TB, and MS conceived and designed the study. ABEG, YM, TB, FW, MW, SWKZ, CS, SG, PSRK and MS performed experiments. FP and MH provided technical support. ABEG, YM, TB, and MS analyzed and interpreted data. MS supervised the project. ABEG, YM, and MS wrote the manuscript. All authors discussed the results and commented on the manuscript.

## Declaration

Authors declare no competing interests.

## Materials and Methods

### Mice and Pig tissues

C57BL/6NRj (aged 8-10 weeks) mice were procured from Janvier Laboratories. All mice were housed on a 12-hour light/12-hour dark cycle with free access to food and drinking water in specific pathogen-free animal facilities. Liver tissues from Landrace pigs were kindly provided by Prof. Constanze Schmidt, University Hospital Heidelberg, Heidelberg University, Heidelberg, Germany (ethics vote – T-19/24). All animal experiments were per the governmental (Regierungspräsidium Karlsruhe, Germany) and institutional (Heidelberg University, Mannheim, Germany) guidelines for the care and ethical use of laboratory animals.

### Liver tissue digestion and subsequent enrichment of LSECs

C57BL/6NRj mice were euthanized by cervical dislocation, and liver tissues were excised for isolating LSECs. The step-by-step method for isolating LSECs is described in the results section. Here, we describe all materials used and the composition of various buffers.

Chopped liver tissue from each mouse were digested in 5 ml liberase-based digestion mix (5 ml of DMEM [Thermo Fisher Scientific, Cat #61965059] + 1 mg Liberase^TM^ Research Grade [Roche, Cat #5401127001] + 0.1 mg DNAse I [Sigma-Aldrich, Cat #10104159001]).

For magnetic enrichment of LSECs, NPCs from each liver were incubated in 200 μl of CD146- selection mix (175 μl FACS buffer + 25 μl of CD146 microbeads [Milentyi Biotec, Cat #130-092- 007]). FACS buffer used for preparing the selection mix and washing the LS columns [Milentyi Biotec, Cat #130-042-401] contains 3% FCS in PBS solution.

For pig liver samples, digested single cell suspension was directly stained with APC-conjugated anti-pig CD31 [BioRad, Cat #MCA1746APC] antibody.

### Preclinical MASH model

MASH was induced in mice by feeding *ad libitum* with either a standard diet [Ssniff, Cat #V1534-000] or Choline deficient L-amino acid-defined diet (CDAA) [Ssniff, Cat # E15666-94] for 10 weeks as previously described^18^. Liver tissue sections were analyzed with Picrosirius red staining to evaluate liver fibrosis as previously described^13^.

### Immunofluorescence staining and flow cytometry

Subsequent to magnetic selection, enriched LSECs were stained for various surface markers [CD31 (MEC 13.3), CD146 (ME9F1), and CD117 (2B8)]. All antibodies were purchased from BioLegend. Dead cells were excluded by FxCycle Violet staining [Thermo Fisher Scientific, Cat #F10347]. Stained cells were analyzed on either BD FACS Melody^TM^ or BD Aria II cell sorting platforms. Total LSECs per liver were calculated using CountBright plus counting beads [Thermo Fisher Scientific, Cat #C36995]. FACS data were analyzed using FlowJo software.

### Immunofluorescence staining on cultured LSECs

Coverslips were placed in 24 well plates and coated with Collagen IV [R&D Systems, Cat #3410- 010-02]. 2 million CD146-enriched LSECs were thereupon added to each well and cultured at 37°C and 5% CO2 for 24 hours in complete mouse endothelial cell medium [Pelo Biotech, Cat #PB-M1168]. Cells were subsequently washed with PBS, fixed in 4% PFA [Carl Roth, Cat #0335.1], blocked with 10% normal donkey serum [Biozol, Cat #LIN-END9010-10] and incubated in a primary antibody mix consisting of CD31 [Dianova, Cat #DIA-310] and CD32b [R&D Systems, Cat #AF1460] overnight. Ensuing washing in PBS, enriched LSEC were incubated with a mix composed of DAPI [Invitrogen, Cat #D1306], AF488-conjugated anti-goat [Dianova, Cat #705-545-147] and Cy3-conjugated anti-rat [Dianova, Cat #712-165-153] secondary antibodies at room temperature and mounted using Dako mounting medium [Agilent, Cat #S302380-2].

### Gene expression analysis

Total RNA from FACS-sorted LSECs was isolated using PicoPure™ RNA isolation kit [Thermo Fisher Scientific, Cat #KIT0204]. Total RNA was transcribed into complementary DNA using a QuantiTect reverse transcription kit [Qiagen, Cat #205313]. Quantitative PCRs (qPCRs) were performed with TaqMan fast advanced master mix [Thermo Fisher Scientific, Cat #4444557]. TaqMan primers (*Actb*, Mm00607939_s1; *ACTB*, Ss03376563_uH; *CDH5*, Ss03378336_u1; *CLDN5*, Ss03373518_u1; *COL4A1*, Ss06915326_mH; *Dll4*, Mm00444619_m1; *Dkk3*, Mm00443800_m1; *Efnb2*, Mm00438670_m1; *Fcgr2b*, Mm00438875_m1; *Il1a*, Mm00439620_m1; *KDR*, Ss03375683_s1; *Kit*, Mm00445212_m1; *Sox17*, Mm00488363_m1; *Stab1*, Mm00460390_m1; *Stab2*, Mm00454684_m1; *Thbd*, Mm00437014_s1; *TIE1*, Ss03373579_g1; and *Wnt9b*, Mm00457102_m1) were ordered from Thermo Fisher Scientific. Gene expression was calculated based on the ΔΔCt relative quantification method. mRNA abundances were normalized to *Actb/ACTB* expression as indicated.

### Statistical analysis

Statistical analysis was performed using GraphPad Prism version 10 (GraphPad Software). Data are expressed as means ± SD. Used statistical tests are indicated in corresponding figure legends. A P value of less than 0.05 was considered statistically significant.

---

**Supplementary Figure S1.**
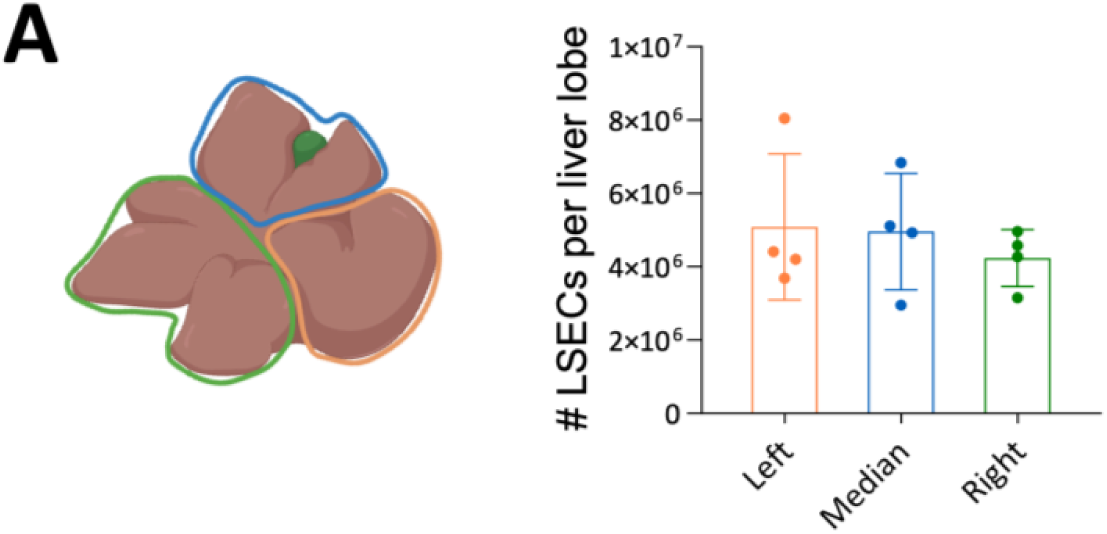
Isolating LSECs from different lobes of the liver. (**A, left**) The liver tissue was divided into three parts as shown in the picture. (**A, right**) Each part was processed to isolate LSECs. The dot plot shows the total count of LSECs isolated from each part of the liver [mean ± SD, n = 4 mice].

**Supplementary Figure S2.**
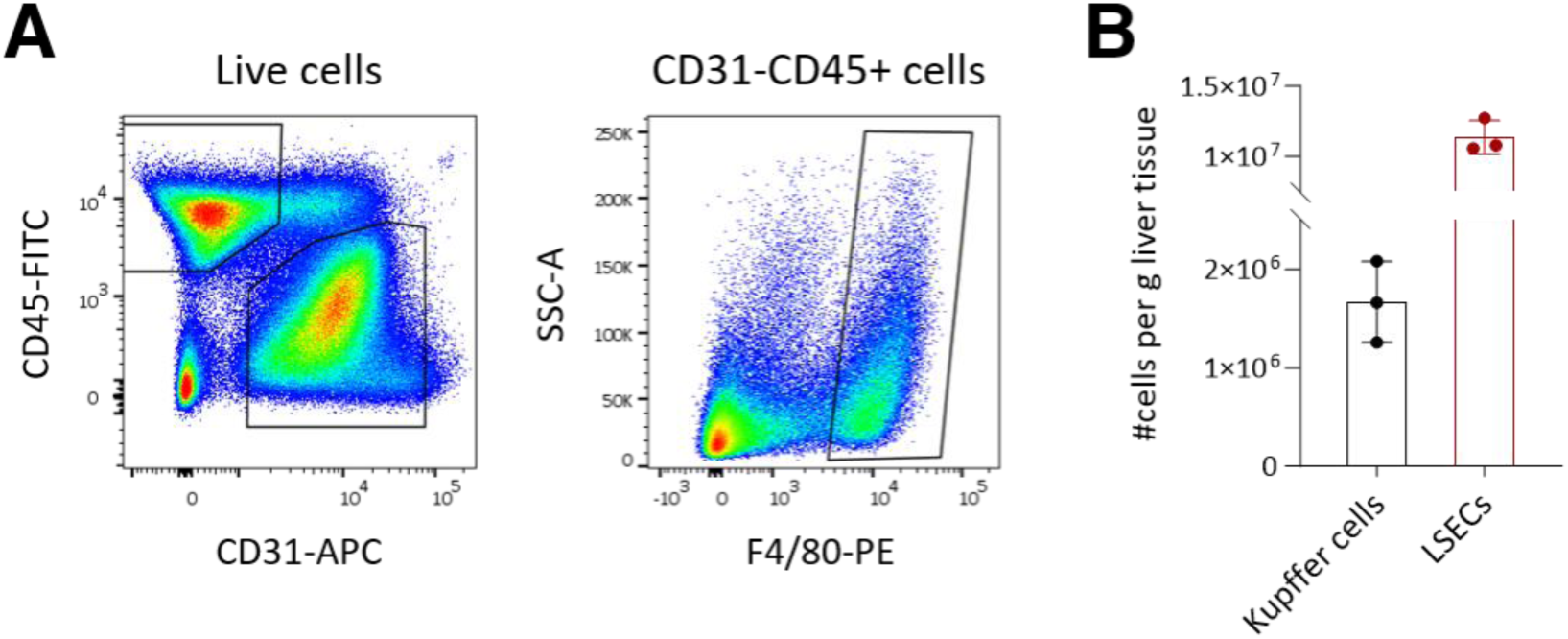
Isolating Kupffer cells and LSECs from the liver. (**A**) FACS gating strategy to analyze and sort LSECs and Kupffer cells. All live cells were stratified based on the expression of CD45 and CD31. CD31+ cells were gated as LSECs. CD31-CD45+ cells were further analyzed for the expression of F4/80. CD31-CD45+F4/80+ cells were isolated as Kupffer cells. (**B**) The dot plot shows the number of isolated LSECs and Kupffer cells from the liver. [mean ± SD, n = 3 mice].

